# Optimized CRISPR/Cas9 Electroporation and Single Cell Cloning Protocol for Generating Pure Cellular Models in Human Immortalized Myoblasts

**DOI:** 10.1101/2025.09.08.674725

**Authors:** Adrien Rihoux, Alexie Gagné, Jean Mezreani, Clémence Gonthier-Cummings, Laura K. Hamilton, Eric Samarut, Martine Tétreault

## Abstract

**Background:** Genome editing in human skeletal muscle research requires protocols that maximize delivery while preserving viability and clonal outgrowth. We sought to develop a reagent-free workflow for CRISPR/Cas9 editing in human immortalized myoblasts and to demonstrate its performance in two use cases, an *IARS1* knockout and an *MLIP* homozygous knock-in.

**Methods:** We optimized electroporation parameters using a green fluorescent protein reporter to compare three electrical settings for transfection and survival in E6/E7 myoblasts, then applied ribonucleoprotein delivery for editing. We evaluated the effect of confluency at electroporation, performed single-cell cloning without antibiotics or fluorescence-activated sorting, and validated edits by high-resolution melting pre-screen followed by Sanger sequencing.

**Results:** Electroporation optimization identified one parameter set that maximized delivery while preserving viability. Performing electroporation at low confluency increased clonal outgrowth and editing rates. The workflow yielded an 84% success rate for *IARS1* knockout and a 3.3% success rate for *MLIP* homozygous knock-in. High-resolution melting provided a very sensitive pre-screen, detecting 96% to 100% of actual edits, reducing the number of Sanger sequencing needed. Performance was reproducible across runs and myoblast lines and increasing single-cell seeding scaled yields without compromising purity.

**Conclusions:** This work provides a practical and reproducible selection-free protocol that couples electroporation optimization, low confluency editing, single-cell cloning, and high-resolution melting sorting to generate pure edited myoblast lines. The approach is applicable to disease modeling in neuromuscular research and clarifies feasibility boundaries for essential genes and homology-directed repair in these cells.

## Background

Genome editing with CRISPR/Cas9 has reshaped biomedical research by enabling precise, targeted modifications in mammalian genomes (1, 2). In neuromuscular disorders, where effective treatments remain scarce, CRISPR allows the recreation or correction of patient variants in human skeletal muscle cells (3). Human immortalized myoblasts provide a particularly valuable model for these applications. They expand robustly in culture and preserve their myogenic identity, ensuring the maintenance of key skeletal muscle characteristics (4, 5). This makes them a reproducible system for investigating disease-associated genetic variants in a human muscle context.

Despite these advantages, editing immortalized human myoblasts requires careful optimization. Myogenic cells are relatively refractory to non viral transfection, and electroporation settings strongly trade off delivery with survival, which determines whether single cells can expand into clones (6, 7). Electroporation supports efficient delivery of Cas9 ribonucleoprotein (RNP) complexes in immortalized human myoblasts, enabling editing and recovery of single cell clones (8, 9). Still, parameters such as voltage, pulse width, and pulse number must be optimized for each line, as they directly influence both uptake and cell death (6, 8, 10). Work with Neon electroporation of Cas9 plasmids in myoblasts highlights the importance of adjusting pulse settings to outperform lipid-based delivery (11). Culture conditions further shape outcomes: lower confluency has been shown to enhance transfection and editing in primary human myoblasts, and this practice is often extended to immortalized lines. However, direct quantitative studies in these models remain limited (7).

A common strategy to enrich edited cells is the use of antibiotic selection or fluorescence-based sorting (12). However, both approaches introduce external pressures that can interfere with the subsequent differentiation of myoblasts as well as downstream analysis. Sorting has been shown to induce oxidative stress and broad shifts in the cellular metabolome that persist into downstream assays (13, 14). Antibiotics likewise alter cell state; penicillin–streptomycin modifies gene expression programs, and streptomycin reduces protein synthesis and myotube differentiation in C2C12 cells (15, 16). To avoid these confounding effects, single-cell cloning provides a cleaner route. Under optimized culture conditions, immortalized human myoblasts can expand from individual cells while maintaining myogenic markers and fusion potential, although clonogenic efficiency differs across lines and between clones (17, 18).

Validation of edited clones must also be rigorous. Amplicon sequencing is the standard approach, but on-target events can be complex and complicate straightforward interpretation (19–21). High-resolution melting (HRM) analysis offers a rapid pre-screen for small sequence changes and indels (22–24). This method allows early triage and reduces the number of clones that must undergo full sequencing (22, 24). HRM can also directly detect CRISPR-induced indels (25), enabling efficient identification while maintaining a chemical-free, single-cell cloning workflow.

In the context of neuromuscular disease diagnosis, next-generation sequencing (NGS) has become the first-tier investigation (26). However, NGS often result in the identification of variants of unknown significance or in novel or newly discovered genes, which require further investigation to confirm functional impact (27). CRISPR/Cas9 gene editing in immortalized and primary myoblasts can serve this purpose, but most protocols have targeted well-described myogenic regulators (7) or highly expressed genes involved in neuromuscular diseases, such as dystrophin (11). Thus, it is essential to develop gene editing workflows that will address the challenges of precision medicine, such as testing the variant itself and/or targeting genes with variable expression levels in muscle or that have a function essential to cell survival.

In this study, we present a selection-free CRISPR/Cas9 editing protocol for immortalized human myoblasts. The workflow integrates optimized electroporation conditions, single-cell cloning without antibiotics or fluorescence sorting, and HRM pre-screening for rapid validation. We demonstrate its utility in two challenging contexts, a knockout of the essential *IARS1* gene and homozygous knock-in of a disease-associated *MLIP* variant, providing a reproducible, step-by-step protocol for generating pure cellular models of neuromuscular disease.

## Methods

### Human immortalized myoblast culture

Two immortalized human myoblast lines were used: E6/E7 (provided by the Montreal Neurological Institute) and 13B13 (provided by Université Laval). Cells were maintained in Sk-Max medium (Wisent, Cat. 301-061-CL) supplemented with Sk-Max growth supplement (Wisent, Cat. 301-061-XL) and 20% fetal bovine serum (FBS; Wisent, Cat. 080-150). Cultures were grown in 100 mm tissue culture dishes (Sarstedt, Cat. 83.3902) at 37°C in a humidified incubator with 5% CO₂ (CellXpert 170i, Eppendorf). Medium was replaced every 48h to prevent acidification, and cultures were monitored by phase-contrast microscopy. Cells were passaged every 2–3 days when reaching ∼70% confluency. For passaging, medium was aspirated, cells rinsed with 2 mL Dulbecco’s phosphate-buffered saline (D-PBS; Wisent, Cat. 311-430-CL) and detached with 2 mL of 0.05% trypsin-EDTA (Wisent, Cat. 325-042-CL) for 3 min at 37°C. The reaction was quenched by adding 8 mL of fresh medium, and cells were reseeded at a 1:5 dilution in new dishes until experimental use.

### Design of *MLIP* knock-in using HDR template

Custom gRNAs and single-stranded HDR templates were designed using online tools (ThermoFisher, Benchling, Integrated DNA Technologies, and Synthego). Selection criteria included high on-target scores, minimal predicted off-target activity, and cut site proximity (<10 bp) to the intended edit. The chosen gRNA targeted *MLIP* at c.2284C>T, introducing a premature stop codon (p.Gln762Ter). The HDR template carried two substitutions: (i) CAG→TAG to generate the nonsense mutation, and (ii) TGG→TCG to disrupt the PAM sequence and prevent re-cleavage after repair.

MLIP gRNA sequence: 5’-TTGGAACAGAAGGTCAGTGT-3’. HDR template sequence: 5’-TAAAAATCACATTAAAAATGTAAAATTCACTGTTTTTGAGCGAACACTGACCTTCTATTCAAAAGCAATGTGTTTGTAGGGATTGCTGCAAAAGCC-3’.

### Design of *IARS1* knockout using Non-Homologous End Joining (NHEJ)

For *IARS1* disruption, a validated TrueGuide™ synthetic gRNA (ThermoFisher, Assay ID CRISPR1056093, Cat. A35533) was used. This gRNA targets exon 3 of *IARS1* (Chr.9: 92288232 – 92288254, GRCh38) with the sequence 5’-GCAACTGGACTGCCTCACTA-3’ (PAM TGG, forward strand). Frameshift-inducing indels generated by NHEJ were expected to disrupt protein function.

### Cas9 nuclease and RNP assembly

Both *MLIP* and *IARS1* editing experiments employed Alt-R™ S.p. Cas9 Nuclease V3 (IDT, Cat. 1081058). gRNAs and Cas9 were assembled into RNP complexes immediately before electroporation to maximize editing efficiency and minimize off-target activity.

### Optimization of Electroporation Parameters

To identify conditions that maximize transfection efficiency while preserving viability, electroporation was performed using the Neon electroporation system (Thermo Fisher Scientific) with the 10 µL tip and chamber. E6/E7 human immortalized myoblasts were electroporated under three protocols varying in voltage, pulse duration, and pulse number: 1475 V with two pulses of 20 ms (protocol 1), 1650 V with three pulses of 10 ms (protocol 2), and 1680 V with a single pulse of 20 ms (protocol 3). For each reaction, 100,000 cells were resuspended in Buffer R (Neon kit) together with 500 ng of GFP-expressing plasmid pmEGFP-C1 (Addgene plasmid #36312) in a total volume of 10 µL. Immediately after electroporation, cells were transferred into 24-well plates (Sarstedt, Cat. 83.3922) containing pre-warmed growth medium and maintained at 37°C in a humidified atmosphere of 5% CO₂. For the 13B13 myoblast line, electroporation was carried out using parameters recommended by Université Laval (1100 V, two pulses of 20 ms).

### Assessment of Electroporation Performance

Twenty-four hours post electroporation, to allow recovery and reporter expression, transfection efficiency and cell viability were assessed by fluorescence microscopy. Images were acquired on an Axio Observer Z1 inverted microscope (ZEISS) equipped with an EC Plan Neofluar 10x/0.30 objective and a onefold TIRF camera adapter. Fluorescence was captured using the 38 HE GFP filter set (excitation 450–490 nm, emission 500–550 nm) with an X-Cite 120LED light source set to 25% intensity, exposure time 800 ms. Bright field images were acquired using transmitted light LED at 18.3% intensity with a 100 ms exposure. Images were recorded with an AxioCam MR R3 camera at 12-bit depth, pixel size 0.645 µm, under ZEISS ZEN. Scale bars were calibrated from pixel dimensions.

Cells electroporated without GFP plasmid served as negative controls for autofluorescence. For each condition, three non-overlapping fields of view were imaged within a single well, sampled at random locations. Viability was defined as the number of adherent cells per field and transfection efficiency as the fraction of GFP positive cells. Viability and transfection efficiency were analyzed as described in data analysis and statistics.

### Electroporation experiment

To apply the optimized parameters to genomic editing, electroporations were carried out using the Neon system (Thermo Fisher Scientific) with the 10 µL chamber. Cells were harvested at approximately 50% confluency, washed twice with PBS, and resuspended in Buffer R. For each reaction, 100,000 cells were combined with the editing reagents in a final volume of 10 µL and loaded into the Neon tip, taking care to avoid air bubbles.

For *MLIP* knock-in experiments, the mixture contained 2.4 µg Alt-R™ S.p. Cas9 Nuclease V3 (IDT), 480 ng gRNA, and 1.2 µL of 100 µM HDR template (IDT). For *IARS1* knockout experiments, the reaction contained Cas9 nuclease and gRNA only, with Buffer R used in place of the HDR template. Immediately after electroporation, cells were transferred into 24-well plates (Sarstedt, Cat. 83.3922) containing 1 mL of pre-warmed culture medium and maintained at 37°C in 5% CO₂. Following four days of recovery, cells were distributed for single-cell cloning as described below.

### Single cell cloning

To generate homogeneous populations of edited cells, single-cell cloning was performed. Cells were diluted to a final concentration of ∼0.8 cells/well to minimize the probability of more than one cell per well, and 150 μL of suspension was dispensed into sterile 96-well plates (Sarstedt, Cat. 83.3924) using a multichannel pipette. Plates were incubated at 37°C with 5% CO₂, and clonal outgrowth was assessed by phase-contrast microscopy after 7 days.

### Clonal expansion

To scale up successfully edited clones for downstream use, colonies were monitored daily by microscopy. After 7–10 days, once clones reached ∼50–60% confluency, they were transferred into progressively larger culture vessels. During the transfer from 48-well to 24-well plates, each clone was divided into two wells: one allocated for genomic DNA extraction and the other for expansion. Edited clones confirmed by molecular analysis were further expanded into 100 mm dishes for downstream applications and cryopreservation.

### Genomic DNA extraction

Genomic DNA was extracted using the PureLink™ Genomic DNA Kit (Invitrogen, Cat. K182002) according to the manufacturer’s protocol, with a final elution volume of 50 µL. DNA was quantified by spectrophotometry and stored at 4°C for short-term use or –20°C for long-term storage.

### High-resolution melting (HRM) analysis

To pre-screen clones prior to sequencing, extracted genomic DNA from candidate clones was analyzed by HRM. Amplicons were designed to span the CRISPR target with lengths of approximately 100 to 200 base pairs to maximize melt-curve resolution near the edit site (sequences listed in Table S1). Each reaction contained 2 ng genomic DNA, 0.5 μL forward primer, 0.5 μL reverse primer, and 5 μL Precision Melt Supermix (Bio-Rad, 1725112), brought to 13 μL with nuclease-free water (Life Technologies, AM9937). Amplification and melting were performed on a Roche LightCycler 96 using white 96-well plates (Progene, 87-C96-LC480-WP). One reaction was run per clone; the same wild-type DNA sample was included in every run as a control.

Cycling conditions were: 95°C for 120 s; 45 cycles of 95°C for 10 s, 60°C for 30 s, and 72°C for 30 s; then high-resolution melting from 65°C to 95°C with continuous acquisitions at 0.2°C increments; and cooling at 37°C for 30 s. Melting profiles were analyzed in LightCycler software.

### Sanger sequencing

Clones identified by HRM as potentially edited were validated by Sanger sequencing. Targeted loci were amplified by PCR in 25 µL reactions containing 25 ng genomic DNA, 0.5 µL forward primer, 0.5 µL reverse primer (sequences listed in Table S1), 0.25 µL Taq DNA polymerase (Froggabio, Cat. TAQP500), 2.5 µL 10x buffer, and 2 µL dNTP mix, with nuclease-free water added to volume. PCR was performed on a ProFlex™ 96-well system (Thermo Fisher Scientific) using the following cycling conditions: initial denaturation at 94°C for 3 min; 35 cycles of 94°C for 30 s, 60°C for 30 s, and 72°C for 1 min; final extension at 72°C for 10 min; indefinite hold at 4°C. Amplicon sizes were 516 bp for MLIP and 636 bp for IARS1.

PCR products were submitted directly (without purification) to Genome Québec (Centre d’expertise et de services Génome Québec, CESGQ) for sequencing using forward primers only at 5 µM. One PCR product was sequenced per clone.

### Chromatogram analysis

Sanger sequencing chromatograms were analyzed with the ICE CRISPR Analysis Tool (Synthego; https://www.synthego.com/products/bioinformatics/analysis). ICE was used to decode mixed traces, estimate indel frequencies, and calculate KO or KI editing scores for each clone.

### Cryopreservation of clones

Validated clones cultured in 100 mm dishes were harvested with 0.05% trypsin-EDTA, pelleted, and resuspended in Human Skeletal Muscle Myoblast Freezing Medium (Celprogen, Cat. 36048-18). Cell suspensions were aliquoted into cryovials and frozen in a controlled-rate freezing container (Nalgene® Mr. Frosty, Thermo Fisher, Cat. 5100-0001) at –80°C for 24–72h, before transfer to liquid nitrogen storage (–196°C).

### Outcome definitions

Growth efficiency was defined as the proportion of single-cell seeded wells that produced at least one clonal outgrowth that could be successfully passaged. Protocol efficiency was defined as the proportion of seeded clones that displayed the intended DNA modification confirmed by Sanger sequencing, relative to the total number of seeded clones. Editing efficiency was defined as the proportion of clones with the intended DNA modification (confirmed by Sanger sequencing) among those that successfully expanded to the point of sequencing.

### Data analysis and statistics

All statistical analyses were performed using GraphPad Prism (version 9.0.0). For electroporation optimization experiments, viability (cell counts per field) and transfection efficiency (GFP-positive ratio) were analyzed by one-way ANOVA. Viability comparisons were made against Healthy cells using Dunnett’s multiple comparison test, with an additional comparison of Protocol 2 versus Control to evaluate plasmid-associated toxicity. Differences in GFP-positive ratios across Protocols 1–3 were assessed with Tukey’s all-pairwise test. Assumptions of normality (Shapiro– Wilk) and homogeneity of variance (Brown–Forsythe and Bartlett) were confirmed prior to ANOVA.

For clone-level categorical outcomes (growth efficiency, editing efficiency, and overall protocol efficiency), results were summarized in contingency tables. Chi-square tests were applied when expected counts were ≥5; otherwise, Fisher’s exact test was used.

HRM assay performance was evaluated against Sanger sequencing by calculating sensitivity, specificity, positive predictive value (PPV), and negative predictive value (NPV).

Graphs display mean ± SD with individual data points overlaid where appropriate. A significance threshold of α = 0.05 was applied throughout. Significance is indicated on figures as *p < 0.05, **p < 0.01, ***p < 0.001, and ****p < 0.0001.

## Results

Testing an editing workflow requires scenarios with distinct biological constraints and direct relevance to neuromuscular disease and precision medicine; thus, we selected two challenging situations to develop our approach. For a stringent loss-of-function challenge, we targeted *IARS1*, an aminoacyl-tRNA synthetase responsible for charging isoleucine to its cognate tRNA. Loss of IARS1 disrupts aminoacylation and halts translation, making the recovery of stable null clones exceptionally difficult (28–30). For a precision benchmark, we pursued a homozygous knock-in of a patient variant in *MLIP*. Biallelic *MLIP* variants are associated with human myopathy characterized by hyperCKemia and rhabdomyolysis, with cardiac involvement reported in some cases (31, 32).

### Electroporation parameters influence cell survivability and transfection efficiency

To determine the best conditions, three electroporation protocols were compared in E6/E7 hiMyo using a GFP reporter plasmid (see methods; Figure 1A). Cell viability differed significantly among groups (one-way ANOVA, F(4,10) = 93.97, p < 0.0001). Relative to non-electroporated control (NEC) cells, viability was reduced in all three protocols (Dunnett adjusted p < 0.0001; Figure 1B). Protocol 2 (1650 V, 3 pulses, 10 ms) yielded the highest survival among electroporated groups. Electroporated control cells (EC), without GFP plasmid, using the same parameters as Protocol 2, displayed viability not significantly different from NEC (adjusted p = 0.0788). When the GFP plasmid was added, viability decreased (Protocol 2 vs EC, adjusted p = 0.0004). The proportion of GFP-positive cells did not differ significantly across Protocols 1–3 (one-way ANOVA, F(2,6) = 0.7103, p = 0.53; all Tukey adjusted p > 0.5; Figure 1C). These data show that electroporation efficacy was similar in all three protocols, but cell survival was higher using protocol 2.

**Figure 1.**
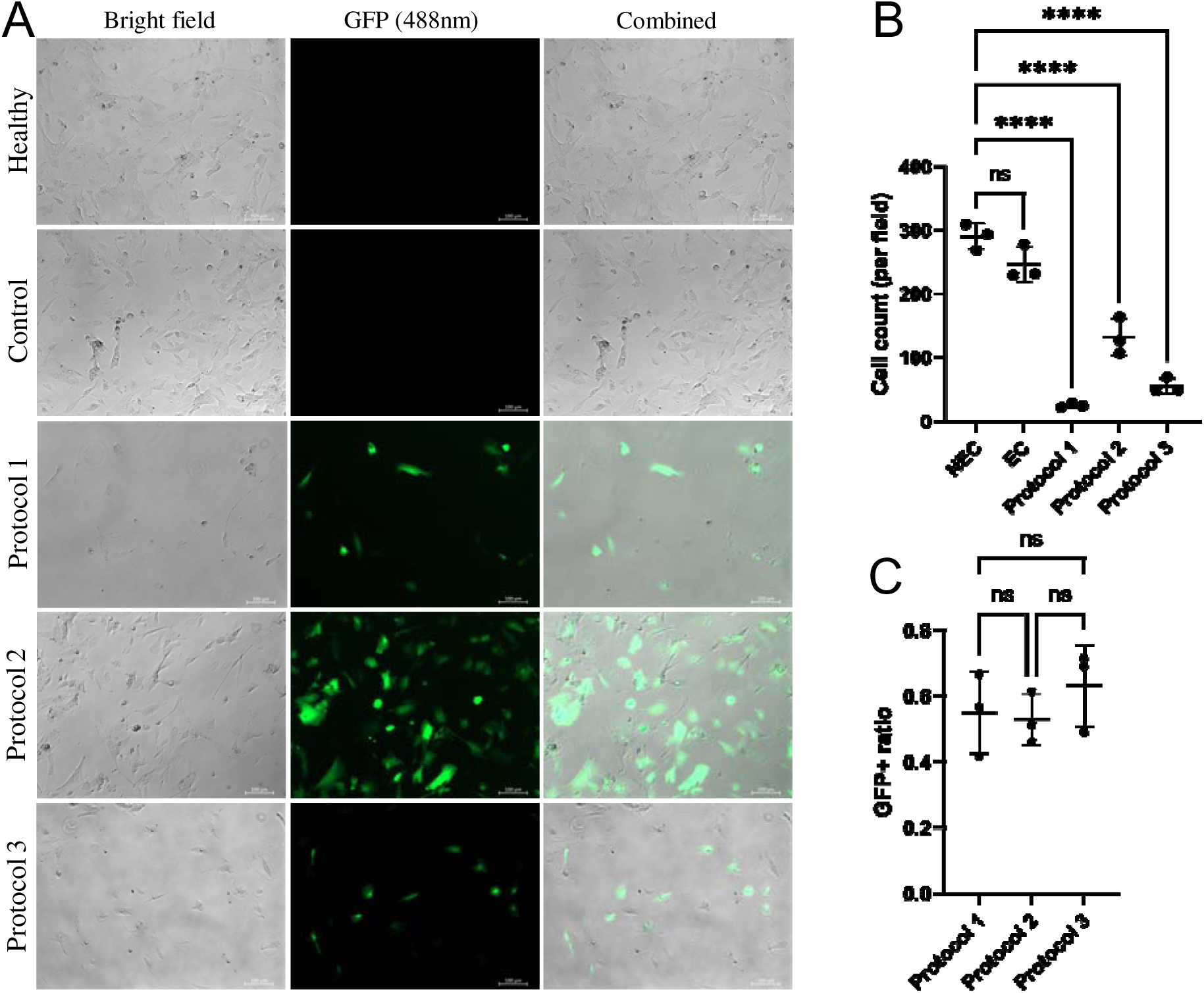
Electroporation optimization in E6/E7 human immortalized myoblasts. (A) Representative bright field, GFP fluorescence, and merge twenty-four hours after electroporation in E6/E7 myoblasts. Conditions displayed in the panels are non-electroporated control (NEC), electroporated control (EC; Protocol 2 electrical settings without GFP plasmid), Protocol 1 (1475 V, 2 pulses, 20 ms), Protocol 2 (1650 V, 3 pulses, 10 ms), and Protocol 3 (1680 V, 1 pulse, 20 ms). Scale bar equals 100 µm. (B) Cell viability as mean ± SD cell counts per field. (C) GFP positive ratio under each condition. Data represent three fields of view within one well per condition.

### Low cell confluency improves growth and protocol efficiency

To assess the impact of confluency, E6/E7 hiMyo were seeded at high (70%) or low (40%) confluency before CRISPR/Cas9-mediated *IARS1* knockout. We first measured cell growth in each condition and observed an increase in growth efficiency from 2.28% at high confluency to 11.00% at low confluency (p < 0.0001; Figure 2A). We then determined the protocol efficiency by measuring the proportion of seeded wells that yielded clones with confirmed edits and observed an increase in the low confluent conditions from 0.72% to 4.56% (p < 0.0001; Figure 2B). Finally, we investigated editing efficiency among surviving clones and again observed an increase from 31.43% to 86.42% (p = 0.0011; Figure 2C) was observed at low confluency. In support to previous reports, our data shows higher growth and editing efficiency at low confluence.

**Figure 2.**
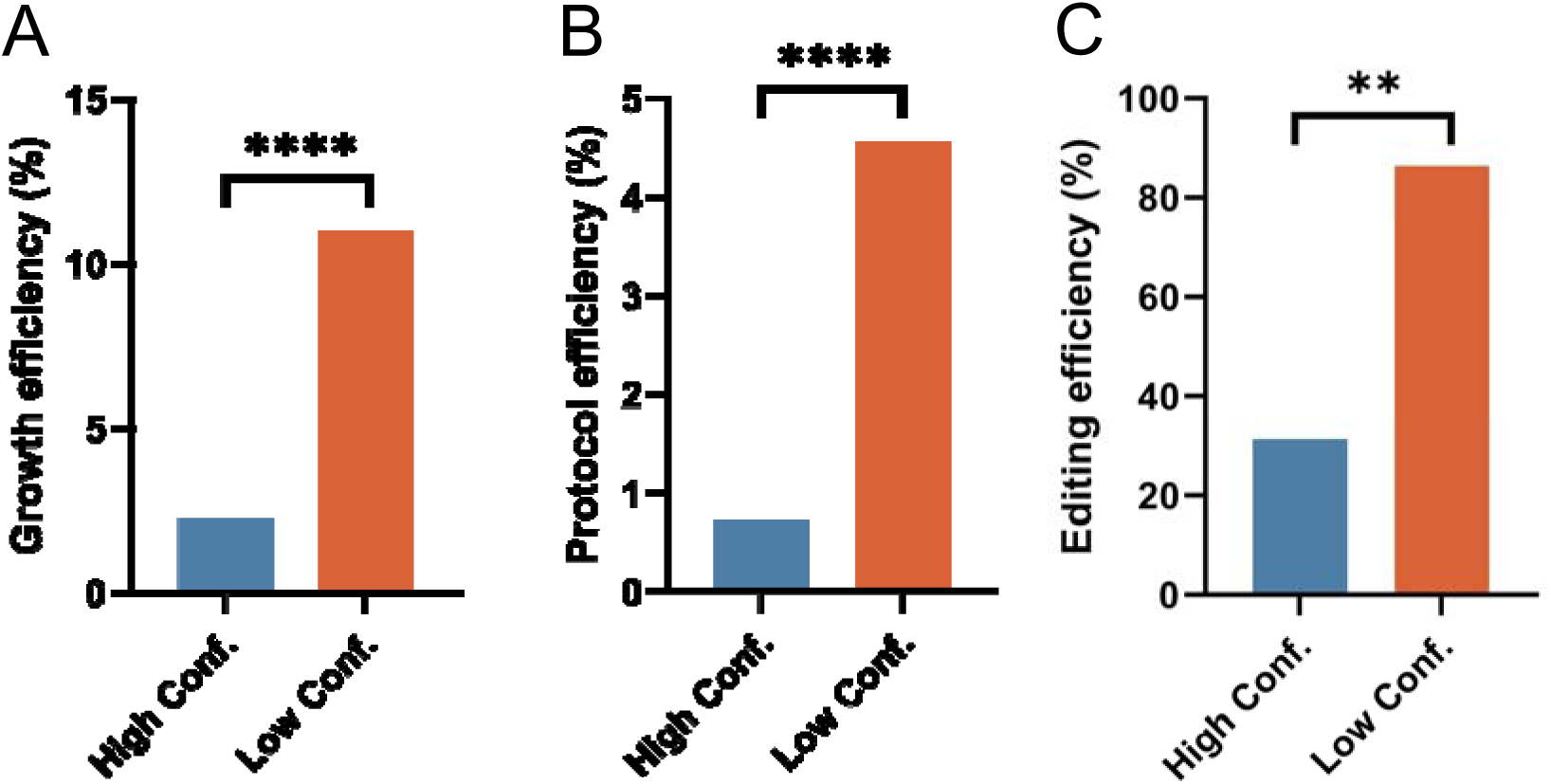
Lower confluency improves growth and editing outcomes in myoblasts. (A) Growth efficiency, proportion of seeded wells that produced viable clonal outgrowths, comparing high (∼70%) and low (∼40%) confluency at electroporation. (B) Protocol efficiency, proportion of seeded wells that yielded clones with confirmed edits by Sanger sequencing. (C) Editing efficiency, proportion of edited clones among sequenced clones. Comparisons were evaluated with Fisher exact or Chi square tests (*p < 0.05; **p < 0.01; ***p < 0.001; ****p < 0.0001).

### HRM assay performance

We confirmed editing of candidate clones using a screening approach combining HRM and Sanger sequencing validation (Figure 3A–C). In *IARS1* knockdown experiments, HRM detected 67 true positives, nine false positives, three false negatives, and two true negatives, corresponding to a sensitivity of 95.71% and specificity of 18.18%. In *MLIP* knock-in experiments (E6/E7 line), HRM detected 105 true positives, four false positives, and no false/true negatives (sensitivity 100%, specificity 0%). In *MLIP* knock-in experiments (13B13 line), HRM detected 47 true positives, 0 false positives or negatives, and 0 true negatives (sensitivity 100%). Positive predictive value (PPV) was 88.16% for *IARS1* and 96.33–100% for *MLIP*. Negative predictive value (NPV) was 40% for *IARS1* and not applicable for *MLIP* (Table 1). An HRM screening, prior to Sanger sequencing, accurately identified positive clones.

**Figure 3.**
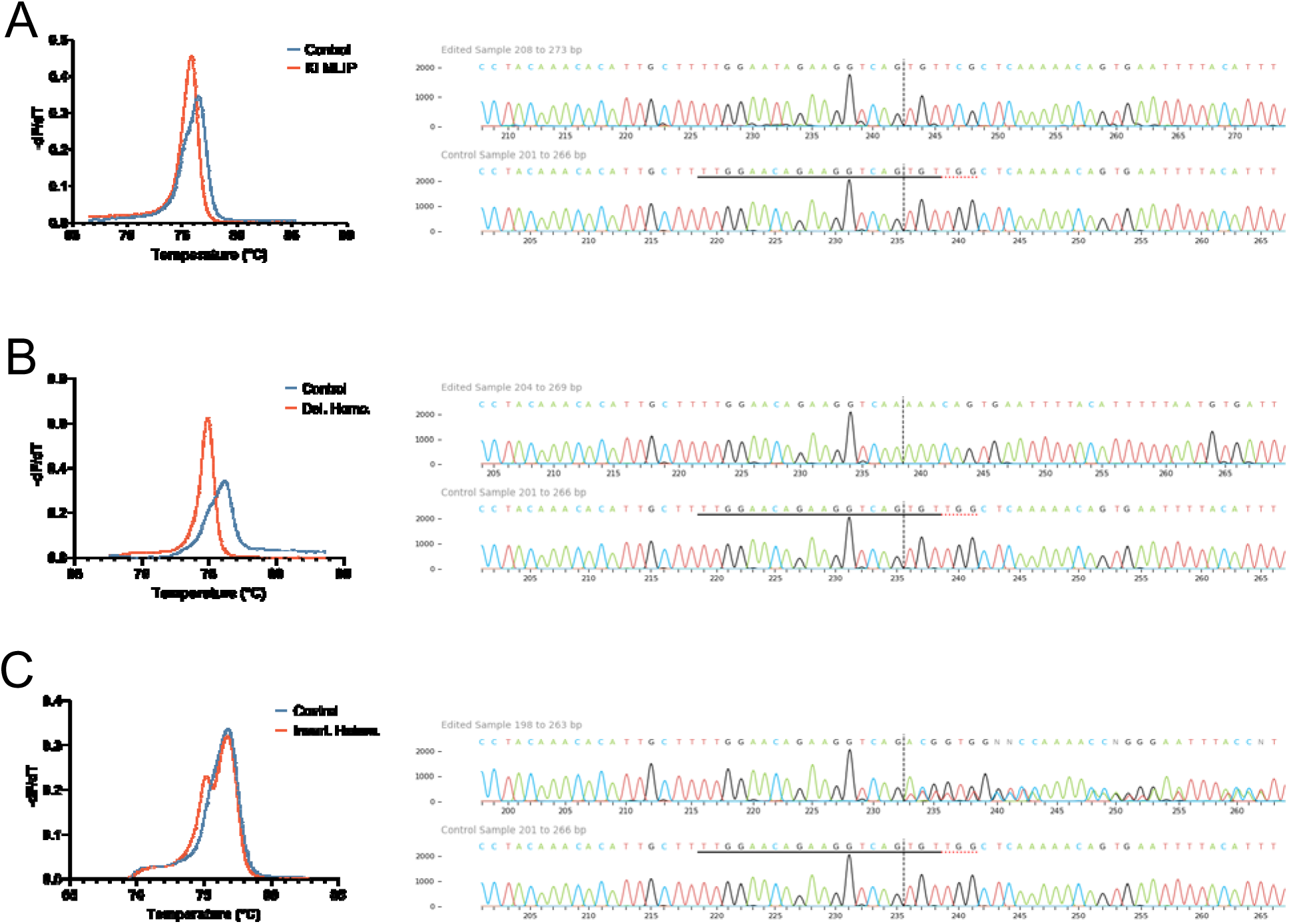
HRM-based detection of CRISPR-induced mutations with Sanger/ICE validation. (A) HRM curve of a homozygous nucleotide substitution compared with the wild type control (left) and corresponding Sanger chromatogram analyzed with the ICE tool (right). (B) HRM curve of a homozygous deletion with its confirmed sequence trace. (C) HRM curve of a heterozygous insertion with corresponding chromatogram. HRM profiles highlight divergence from the wild type, prompting sequencing validation.

**Table 1:**
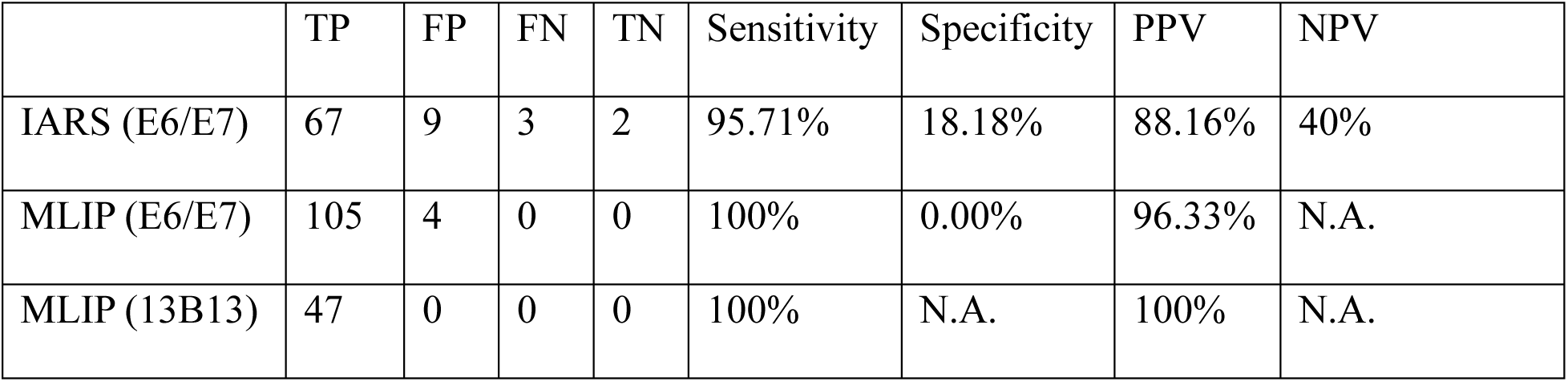
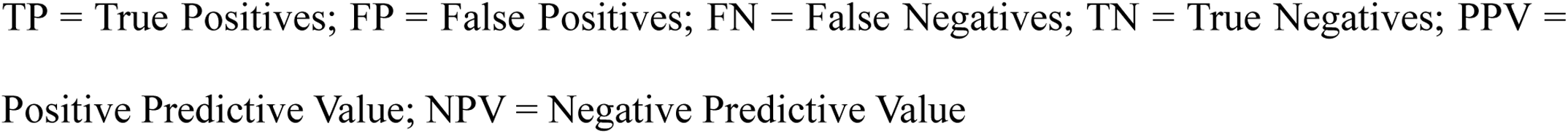
Performance of the high-resolution melting assay in KO IARS and KI MLIP experiments validated by Sanger sequencing.

|  | TP | FP | FN | TN | Sensitivity | Specificity | PPV | NPV |
| --- | --- | --- | --- | --- | --- | --- | --- | --- |
| IARS (E6/E7) | 67 | 9 | 3 | 2 | 95.71% | 18.18% | 88.16% | 40% |
| MLIP (E6/E7) | 105 | 4 | 0 | 0 | 100% | 0.00% | 96.33% | N.A. |
| MLIP (13B13) | 47 | 0 | 0 | 0 | 100% | N.A. | 100% | N.A. |

TP = True Positives; FP = False Positives; FN = False Negatives; TN = True Negatives; PPV = Positive Predictive Value; NPV = Negative Predictive Value

### Reproducibility of electroporation experiments

Two independent *MLIP* knock-in experiments were performed in different myoblast lines (#1: E6/E7 and #2: 13B13). To evaluate the reproducibility of our protocol, we compared growth, protocol and editing efficiencies. Growth efficiency was 22.33% and 19.66% (p = 0.2348; Figure 4A). Protocol efficiency was 0.26% in both experiments (p > 0.9999; Figure 4B), while editing efficiency was 3.27% and 4.17% (p = 0.6739; Figure 4C). None of these comparisons reached statistical significance, which indicates similar performances in both cell lines. To increase the editing efficacy, we performed the gene editing in KI MLIP #2 with an HDR enhancer, but it did not yield a significant improvement (Figure 4A-C). For the KI assays, only clones confirmed by Sanger sequencing to carry the specific homozygous edit were included in editing efficiency calculations, which accounts for the relatively low efficiency values reported.

**Figure 4.**
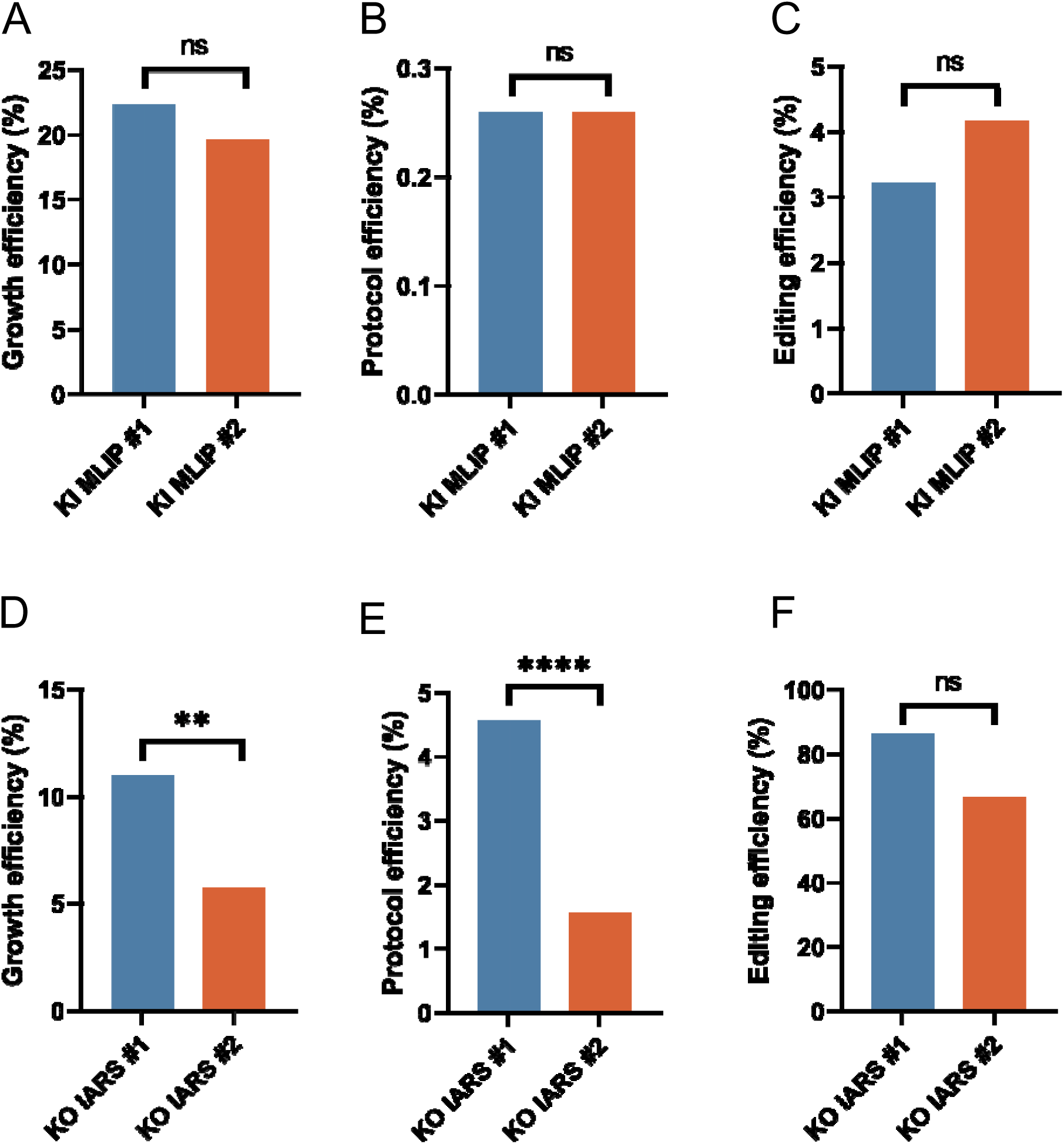
Reproducibility of the electroporation protocol in MLIP knock-in and IARS1 knockdown experiments. (A) Growth efficiency of MLIP knock-in in E6/E7 and 13B13 myoblasts. (B) Protocol efficiency for the same experiments. (C) Editing efficiency, showing similar outcomes between lines. The second run included HDR enhancer (see Methods). (D) Growth efficiency of IARS1 knockout with 1536 versus 384 seeded cells. (E) Protocol efficiency under the same conditions. (F) Editing efficiency, similar across seeding densities.

Two *IARS1* knockdown experiments were performed with different numbers of cells seeded for single-cell cloning (#1: 1536 vs. #2: 384). As for the KI models, we measured the different efficiency parameters. Growth efficiency was 11.00% versus 5.73% (p = 0.0046; Figure 4D), and protocol efficiency was 4.56% versus 1.56% (p = 0.0092; Figure 4E), both statistically significant. Editing efficiency was 86.42% versus 66.67% (p = 0.1428; Figure 4F), which did not differ significantly. These results support a higher growth and protocol efficiency with more cells seeded. A complete overview of raw clone counts and derived efficiencies for each experiment is provided in Supplementary Table S2. Taken together, across all experiments, the overall efficiency was 3.26% for *MLIP* knock-in and 84.4% for *IARS1* knockdown.

## Discussion

This study aimed to develop and validate a reproducible CRISPR/Cas9 protocol tailored to human immortalized myoblasts. We deliberately evaluated the workflow under two stringent conditions representing concrete examples of precision medicine: disruption of *IARS1*, an essential gene whose complete knockout is expected to be incompatible with survival, and homozygous knock-in of a pathogenic *MLIP* variant, a technically demanding application of homology-directed repair (HDR). By testing the protocol in both a loss-of-function scenario and a precise knock-in, we demonstrate its robustness while highlighting the biological and technical boundaries inherent to genome editing in myoblasts.

Optimization of electroporation parameters proved critical for balancing delivery efficiency with cell survival. We found that viability varied significantly across protocols, with Protocol 2 (1650 V, 3 pulses, 10 ms) providing the best compromise between efficiency and survival in E6/E7 cells. This finding aligns with reports that electroporation conditions are cell line specific and must be tuned empirically (6, 8, 10). We also demonstrated that without GFP, electroporated control (using protocol 2) and NEC did not differ significantly in terms of survivability, suggesting a potential toxicity from the GFP in our cell line. An additional determinant of outcome was cell density at the time of electroporation. Low confluency (40%) produced markedly higher growth efficiency, protocol efficiency, and editing efficiency compared to high confluency (70%). These results extend previous work in primary myoblasts, where lower confluency enhances proliferation and editing, to immortalized lines, supporting the principle that reduced confluency favors the recovery of edited clones (7). This observation is critical in the context of myoblast studies, since most gene editing protocols recommend high cell confluence.

Our comparison of KO and KI outcomes further illustrates the influence of repair pathways on editing efficiency. NHEJ, the predominant repair mechanism in mammalian cells, enabled efficient *IARS1* disruption, with an overall editing efficiency of 84.40%. In contrast, HDR-dependent *MLIP* KI reached only 3.26% efficiency. This discrepancy reflects the intrinsic bias of mammalian DNA repair. NHEJ is active throughout the cell cycle, whereas HDR is largely confined to the late S and G2 phases when sister chromatids are available (33, 34). Consequently, HDR-based knock-in is inherently less frequent than NHEJ-mediated editing. Experimental strategies that synchronize cells in S/G2 or restrict Cas9 activity to these phases, as well as the use of small-molecule HDR enhancers, have been shown to increase knock-in efficiency across multiple cell types (35, 36). In our case, the use of HDR enhancers did not improve *MLIP* editing efficiency.

The choice of the targeted gene also influenced the protocol efficiency. *IARS1*, an essential aminoacyl tRNA synthetase, exemplifies the limitations of editing genes critical for basic cellular function. Recovery of complete knockouts was not possible, and only clones carrying partial loss-of-function alleles could be expanded. This outcome is consistent with findings in other models where disruption of essential gene compromises viability (37). In contrast, *MLIP* knock-in, although less efficient, yielded viable clones carrying homozygous edits. Together, these results illustrate that the biological role of the target gene must be carefully considered when designing editing experiments in myoblasts.

Single-cell cloning represented another major constraint. Myoblasts are sensitive to stress during isolation and generally perform better under conditions that minimize disruption of their microenvironment (38, 39). These properties contribute to stochastic variability in survival across experiments, as observed in our *IARS1* knockout assays, where differences in the number of cells seeded significantly influenced growth and protocol efficiency. While such variability complicates the prediction of absolute success rates, single-cell cloning remains essential for generating pure edited lines. Importantly, our approach avoided antibiotics or fluorescence-activated cell sorting, both of which can impose additional stress, alter metabolic states, affect differentiation potential or impact downstream analysis, such as gene expression (13–16). This selection-free workflow therefore preserves the integrity of clones and is particularly well-suited for disease modeling applications.

Validation of edited clones relied on HRM as a pre-screen, followed by Sanger sequencing for final validation. HRM demonstrated very high sensitivity (95.7–100%), ensuring that nearly all true edits were detected. Specificity was lower, particularly in *IARS1* assays, where false positives reduced predictive value. In practice, this meant that additional unedited clones were carried forward for sequencing and short-term culture maintenance, as seen in the IARS1 experiments, where 12 supplemental clones had to be expanded (Table 1). Nonetheless, because all clones underwent confirmatory Sanger sequencing, the risk of retaining false positives was eliminated. Practically, HRM reduced the number of clones requiring sequencing by approximately half, thereby conserving culture resources and minimizing the duration of clonal expansion (Tables S2). These features highlight HRM as a cost-effective and minimally invasive complement to sequencing in workflows where large numbers of clones must be triaged.

Although this study has some limitations, the results highlight the practicality and reproducibility of the protocol. HDR efficiency was modest, and the use of an HDR enhancer did not provide a measurable benefit in our hands, indicating that additional strategies such as synchronization or template design could be explored in the future. Likewise, variability in single-cell cloning outcomes highlights the importance of replication and careful adjustment of seeding density. These considerations, however, do not detract from the central outcome: a straightforward and reproducible protocol that can be readily adapted to diverse myoblast lines and editing goals. By demonstrating both a technically demanding homozygous knock-in and an essential gene knockout, this study establishes a robust foundation for researchers initiating CRISPR-based investigations in neuromuscular disease.

## Conclusion

In conclusion, this work provides a simple and reproducible workflow for CRISPR/Cas9 editing in immortalized human myoblasts. Testing a GFP plasmid under low-confluency conditions allows rapid optimization of electroporation parameters, while preliminary single-cell cloning confirms that a given line can sustain clonal expansion. Once these steps are established, HRM offers a rapid pre-screening tool that reduces sequencing workload, and the protocol as a whole, reliably yields pure edited clones without relying on antibiotics or FACS. Importantly, reproducibility was demonstrated in two independent lines and across distinct editing contexts, underscoring its general utility. Beyond these specific results, the protocol offers a practical entry point for investigators new to CRISPR editing in muscle cells and provides a flexible framework for disease modeling in neuromuscular research.

## List of abbreviations

CRISPR: Clustered Regularly Interspaced Short Palindromic Repeats
FACS: Fluorescence-Activated Cell Sorting
GFP: Green Fluorescent Protein
HDR: Homology-Directed Repair
hiMyo: Human immortalized myoblasts
HRM: High-Resolution Melting
IARS1: Isoleucyl-tRNA Synthetase 1
KI: Knock-in
KO: Knock-out
MLIP: Muscular Lamin-Interacting Protein
NHEJ: Non-Homologous End Joining
RNP: Ribonucleoprotein

## Declarations

ES is a co-founder of DanioDesign Inc. (Qc, Canada) and of Osta Therapeutics (France); The commercial affiliations did not play any role in investigational design, data collection and analysis, the decision to publish or the preparation of the manuscript.

## References

1. Jiang F, Doudna JA. CRISPR-Cas9 Structures and Mechanisms. Annu Rev Biophys. 2017;46:505– 29.

2. Long C, Amoasii L, Bassel-Duby R, Olson EN. Genome Editing of Monogenic Neuromuscular Diseases: A Systematic Review. JAMA Neurol. 2016;73(11):1349–55.

3. Young CS, Pyle AD, Spencer MJ. CRISPR for Neuromuscular Disorders: Gene Editing and Beyond. Physiology (Bethesda). 2019;34(5):341–53.

4. Mamchaoui K, Trollet C, Bigot A, Negroni E, Chaouch S, et al. Immortalized pathological human myoblasts: towards a universal tool for the study of neuromuscular disorders. Skeletal Muscle. 2011;1(1):34.

5. Thorley M, Duguez S, Mazza EMC, Valsoni S, Bigot A, et al. Skeletal muscle characteristics are preserved in hTERT/cdk4 human myogenic cell lines. Skeletal Muscle. 2016;6(1):43.

6. Lojk J, Mis K, Pirkmajer S, Pavlin M. siRNA delivery into cultured primary human myoblasts--optimization of electroporation parameters and theoretical analysis. Bioelectromagnetics. 2015;36(8):551– 63.

7. Goullée H, Taylor RL, Forrest ARR, Laing NG, Ravenscroft G, et al. Improved CRISPR/Cas9 gene editing in primary human myoblasts using low confluency cultures on Matrigel. Skeletal Muscle. 2021;11(1):23.

8. Poyatos-García J, Blázquez-Bernal Á, Selva-Giménez M, Bargiela A, Espinosa-Espinosa J, et al. CRISPR-Cas9 editing of a TNPO3 mutation in a muscle cell model of limb-girdle muscular dystrophy type D2. Mol Ther Nucleic Acids. 2023;31:324–38.

9. Lemoine J, Dubois A, Dorval A, Jaber A, Warthi G, et al. Correction of exon 2, exon 2–9 and exons 8–9 duplications in DMD patient myogenic cells by a single CRISPR/Cas9 system. Scientific Reports. 2024;14(1):21238.

10. Zhang S, Shen J, Li D, Cheng Y. Strategies in the delivery of Cas9 ribonucleoprotein for CRISPR/Cas9 genome editing. Theranostics. 2021;11(2):614–48.

11. Pini V, Mariot V, Dumonceaux J, Counsell J, O’Neill HC, et al. Transiently expressed CRISPR/Cas9 induces wild-type dystrophin in vitro in DMD patient myoblasts carrying duplications. Scientific Reports. 2022;12(1):3756.

12. Mikkelsen NS, Bak RO. Enrichment strategies to enhance genome editing. Journal of Biomedical Science. 2023;30(1):51.

13. Llufrio EM, Wang L, Naser FJ, Patti GJ. Sorting cells alters their redox state and cellular metabolome. Redox Biol. 2018;16:381–7.

14. Ryan K, Rose RE, Jones DR, Lopez PA. Sheath fluid impacts the depletion of cellular metabolites in cells afflicted by sorting induced cellular stress (SICS). Cytometry A. 2021;99(9):921–9.

15. Ryu AH, Eckalbar WL, Kreimer A, Yosef N, Ahituv N. Use antibiotics in cell culture with caution: genome-wide identification of antibiotic-induced changes in gene expression and regulation. Sci Rep. 2017;7(1):7533.

16. He C, Eggelbusch M, Huijts JY, Shi A, de Wit GJ, et al. The commonly used antibiotic streptomycin reduces protein synthesis and differentiation in cultured C2C12 myotubes. Physiol Rep. 2025;13(12):e70353.

17. Massenet J, Gitiaux C, Magnan M, Cuvellier S, Hubas A, et al. Derivation and Characterization of Immortalized Human Muscle Satellite Cell Clones from Muscular Dystrophy Patients and Healthy Individuals. Cells. 2020;9(8).

18. Lathuiliere A, Vernet R, Charrier E, Urwyler M, Von Rohr O, et al. Immortalized human myoblast cell lines for the delivery of therapeutic proteins using encapsulated cell technology. Mol Ther Methods Clin Dev. 2022;26:441–58.

19. Leibowitz ML, Papathanasiou S, Doerfler PA, Blaine LJ, Sun L, et al. Chromothripsis as an on-target consequence of CRISPR-Cas9 genome editing. Nat Genet. 2021;53(6):895–905.

20. Kosicki M, Tomberg K, Bradley A. Repair of double-strand breaks induced by CRISPR–Cas9 leads to large deletions and complex rearrangements. Nature Biotechnology. 2018;36(8):765–71.

21. Cullot G, Boutin J, Toutain J, Prat F, Pennamen P, et al. CRISPR-Cas9 genome editing induces megabase-scale chromosomal truncations. Nature Communications. 2019;10(1):1136.

22. Er T-K, Chang J-G. High-resolution melting: Applications in genetic disorders. Clinica Chimica Acta. 2012;414:197–201.

23. Whitford R. Crop Breeding: Methods and Protocols2014.

24. Wittwer CT. High-resolution DNA melting analysis: advancements and limitations. Hum Mutat. 2009;30(6):857–9.

25. Denbow C, Ehivet SC, Okumoto S. High Resolution Melting Temperature Analysis to IdentifyCRISPR/Cas9 Mutants from Arabidopsis. Bio Protoc. 2018;8(14):e2944.

26. Marchant RG, Bryen SJ, Bahlo M, Cairns A, Chao KR, et al. Genome and RNA sequencing boost neuromuscular diagnoses to 62% from 34% with exome sequencing alone. Ann Clin Transl Neurol. 2024;11(5):1250–66.

27. Ma N, Zhang JZ, Itzhaki I, Zhang SL, Chen H, et al. Determining the Pathogenicity of a Genomic Variant of Uncertain Significance Using CRISPR/Cas9 and Human-Induced Pluripotent Stem Cells. Circulation. 2018;138(23):2666–81.

28. Wang T, Birsoy K, Hughes NW, Krupczak KM, Post Y, et al. Identification and characterization of essential genes in the human genome. Science. 2015;350(6264):1096–101.

29. Hart T, Chandrashekhar M, Aregger M, Steinhart Z, Brown KR, et al. High-Resolution CRISPR Screens Reveal Fitness Genes and Genotype-Specific Cancer Liabilities. Cell. 2015;163(6):1515–26.

30. Rubio Gomez MA, Ibba M. Aminoacyl-tRNA synthetases. Rna. 2020;26(8):910–36.

31. Lopes Abath Neto O, Medne L, Donkervoort S, Rodríguez-García ME, Bolduc V, et al. MLIP causes recessive myopathy with rhabdomyolysis, myalgia and baseline elevated serum creatine kinase. Brain. 2021;144(9):2722–31.

32. Salzer-Sheelo L, Fellner A, Orenstein N, Bazak L, Lev-El Halabi N, et al. Biallelic truncating variants in the muscular A-type lamin-interacting protein (MLIP) gene cause myopathy with hyperCKemia. Eur J Neurol. 2022;29(4):1174–80.

33. Lin S, Staahl BT, Alla RK, Doudna JA. Enhanced homology-directed human genome engineering by controlled timing of CRISPR/Cas9 delivery. Elife. 2014;3:e04766.

34. Matsumoto D, Tamamura H, Nomura W. A cell cycle-dependent CRISPR-Cas9 activation system based on an anti-CRISPR protein shows improved genome editing accuracy. Commun Biol. 2020;3(1):601.

35. Devkota S. The road less traveled: strategies to enhance the frequency of homology-directed repair (HDR) for increased efficiency of CRISPR/Cas-mediated transgenesis. BMB Rep. 2018;51(9):437–43.

36. Li G, Yang X, Luo X, Wu Z, Yang H. Modulation of cell cycle increases CRISPR-mediated homology-directed DNA repair. Cell Biosci. 2023;13(1):215.

37. Wang B, Wang Z, Wang D, Zhang B, Ong S-G, et al. krCRISPR: an easy and efficient strategy for generating conditional knockout of essential genes in cells. Journal of Biological Engineering. 2019;13(1):35.

38. Benedetti A, Cera G, De Meo D, Villani C, Bouche M, et al. A novel approach for the isolation and long-term expansion of pure satellite cells based on ice-cold treatment. Skelet Muscle. 2021;11(1):7.

39. Kulesza A, Burdzinska A, Szczepanska I, Zarychta-Wisniewska W, Pajak B, et al. The Mutual Interactions between Mesenchymal Stem Cells and Myoblasts in an Autologous Co-Culture Model. PLOS ONE. 2016;11(8):e0161693.

