## Supplemental Table 1 for "Optimized CRISPR/Cas9 Electroporation and Single Cell Cloning Protocol for Generating Pure Cellular Models in Human Immortalized Myoblasts"

**Table S1. Primer sequences used for HRM and PCR/Sanger validation**

| **Target gene** | **Application** | **Primer name** | **Sequence (5’–3’)** | **Amplicon size** |
| --- | --- | --- | --- | --- |
| **MLIP** | HRM | MLIP_HRM88_F | GCAGCAATCCCTACAAACAC | 88 bp |
|  | HRM | MLIP_HRM88_R | AACAGAAATCTTCCCAACAG |  |
|  | PCR/Sanger | MLIP_GCD_F | GTGGCATTTCTTCGCTTCTC | 516 bp |
|  | PCR/Sanger | MLIP_GCD_R | CCCAAAGTAGCTTGACAGTGG |  |
| **IARS1** | HRM | HRM_IARS_F_204 | TGAAGACTCCACTATGACAAGAGC | 204 bp |
|  | HRM | HRM_IARS_R_204 | AACCCACTCTGGTGAGCATATC |  |
|  | PCR/Sanger | KO_IARS_F | AAGGCAGGACCACCAGTAAAT | 636 bp |
|  | PCR/Sanger | KO_IARS_R | CTCATCACAATTGCTCGGCAC |  |
