## Supplemental Table 2 for "Optimized CRISPR/Cas9 Electroporation and Single Cell Cloning Protocol for Generating Pure Cellular Models in Human Immortalized Myoblasts"

### Table S2. Raw clone counts and calculated efficiencies for each experiment

| Experiment | Cells seeded | Outgrowth clones | Sequenced clones | Edited clones | Growth efficiency (%) | Editing efficiency (%) | Protocol efficiency (%) | Significance |
| --- | --- | --- | --- | --- | --- | --- | --- | --- |
| MLIP KI #1 (E6/E7) | 1536 | 343 | 128 | 4 | 22.33 | 3.23 | 0.26 | ns |
| MLIP KI #2 (13B13) | 768 | 151 | 50 | 2 | 19.66 | 4.17 | 0.26 | ns |
| IARS1 KO #1 (E6/E7) | 1536 | 169 | 81 | 70 | 11.00 | 86.42 | 4.56 | ** |
| IARS1 KO #1 (E6/E7) | 384 | 22 | 9 | 6 | 5.73 | 66.67 | 1.56 | ** |

Legend: ns, not significant; *p < 0.05; **p < 0.01; ***p < 0.001; ****p < 0.0001. Statistical details are provided in the main text and Results section.
